# Differential amino acid usage leads to ubiquitous edge effect in proteomes across domains of life that can be explained by amino acid secondary structure propensities

**DOI:** 10.1101/2024.07.12.599492

**Authors:** Juliano Morimoto, Zuzanna Pietras

**Affiliations:** Institute of Mathematics, School of Natural and Computing Sciences, University of Aberdeen, Fraser Noble Building, Aberdeen, UK AB24 3UE; Programa de Pós-graduação em Ecologia e Conservação, Universidade Federal do Paraná, Curitiba, 82590-300, Brazil; Department of Physics, Chemistry and Biology (IFM), Linköping University, Sweden

**Keywords:** Genetic code, Structural biology, Environmental responses, Physiology

## Abstract

**Background:** Amino acids are the building blocks of proteins and enzymes, which are pivotal for life on Earth. Amino acid usage provides critical insights into the functional constraints acting on proteins and illuminates molecular mechanisms underpinning traits. Despite this, we have limited knowledge of the genome-wide signatures of amino acid usage across domains of life, precluding new genome and proteome patterns to being discovered.

**Results:** Here, we analysed the proteomes of 5,590 species across four domains of life and found that only a small subset of amino acids is most and least frequently used across proteomes. This creates a ubiquitous ‘edge effect’ on amino acid usage diversity by rank that arises from protein secondary structural constrains. This edge effect was not driven by the evolutionary chronology of amino acids, showing that functional rather than evolutionary constrains shape amino acid usage in the proteome. We also tested contemporary hypotheses about similarities in amino acid usage profiles and the relationship between amino acid usage and growth temperature, and found that, contrary to previous beliefs, amino acid usage varies across domains of life and temperature only weakly contributes to variance in amino acid usage.

**Conclusion:** We have described a novel and ubiquitous pattern of amino acid usage signature across genomes, which reveals how structural constrains shape amino acid usage at the proteome level. This can ultimately influence the way in which we probe deep evolutionary relationships of protein families across the tree of life and engineer biology in synthetic biology.

## Introduction

Proteins perform a wide variety of vital roles which depend on their structure and ultimately, their amino acid composition[1, 2]. The frequency of, or changes to, amino acid profiles in the proteome provide insights into evolutionary mechanisms shaping genomes and their products[3–8] and can be used in disease diagnostics[9, 10] and synthetic biology[11–13]. Despite this, we still lack a proper understanding of amino acid usage and frequency across domains of life, and how proteomes change as a result of the environment and intrinsic physiological constrains imposed by evolving organisms.

Two competing hypotheses exist regarding similarities and differences in amino acid usage in proteomes among species. On the one hand, previous studies have shown that amino acid profiles across proteomes differ among domains of life primarily due to lifestyle[4, 5, 14]. On the other hand, more recent studies suggested that proteomes are conserved among species from different domains of life and contain lineage-specific information on the amino acid requirements to improve fitness[15, 16]. This hypothesis is striking because only specific protein sites (e.g. secondary structures in catalytic motifs) are highly conserved, with the remaining of the protein sequence evolving under fewer biophysical and structural constrains[6, 7, 17]. Moreover, not all proteins are highly expressed which is known to slow down protein sequence evolution and increase conservation of sequences[18, 19]. Nonetheless, amino acid usage appears remarkably consistent[15, 16]. A possible justification for why amino acid profiles might be similar across domains of life is that amino acid profiles are shaped by universal energetic and biophysical constrains and the proteome reflects the overall cost minimization of amino acid synthesis, scavenging opportunities and protein production[3, 20–22]. Conflicting evidence for the conservation of amino acid profiles in proteomes across domains of life exist[4, 8, 14, 15, 20, 23, 24], but data is taxonomically limited preventing a proper test of the generality of the hypothesis. Furthermore, other investigations that reveal new patterns in amino acid usage have not been conducted, rendering our understanding of proteome patterns incomplete.

Here, we collated a comprehensive dataset of 5,590 proteomes of species from across four domains of life (bacteria, eukaryote, viruses, and archaea) to test whether amino acid usage profiles differ or not among distant groups. Next, we incorporated optimal growth temperature for 296 of these proteomes to test whether differences in amino acid usage was indeed a product of the interaction with the environment. We then explored our database to uncover novel patterns in amino acid usage across proteomes. In particular, we tested whether amino acid usage shaped amino acid rank order and if these rank order based on usage correlated with the evolutionary origins of amino acids in living organisms. Our findings advance our understanding of proteomes and open new avenues of research on the patterns underlying amino acid usage across domains of life.

## Results and discussion

### Proteomes differ among distant domains of life

Two competing hypotheses suggest that amino acid usage in proteomes differ or not among distant groups of living organisms. To test this, we firstly analysed the proteome of 5,590 species with complete annotated genomes and identified taxonomy from the NCBI database (see Supplementary Materials for Materials and Methods and Extended Data 1). For each species, we calculated the amino acid profile as the sum of individual amino acid frequencies divided by the total sum of amino acid counts as in[15], which resulted in amino acid profiles for 328 Archaea species, 4,107 Bacteria species, 1,118 Eukaryotes and 37 Viruses. In our data (F_1,111792_ = 7247.30, p < 0.001), as in previous studies[3], the number of redundant codons and the frequency for the corresponding amino acid were positively correlated and to account for this, we used standardised amino acid frequency (i.e. amino acid frequency divided by the number of redundant codons). We found no evidence that the amino acid frequencies were conserved across domains of life as shown by both differences in the principal component analysis (PCA) clusters (Fig 1a-b) and in the average amino acid usage frequency across domains (Domain*Amino acid frequency: F_3,111720_ = 398.9, p < 0.001, Fig 1c). This effect was driven by proteome-wide differences in amino acid frequencies and not by a single or few amino acids having disproportionate effect, as shown in our PCA analysis (Fig 1a) and the average amino acid usage frequencies across domains (Fig 1c). Nonetheless, it is worth highlighting that a nearly two-fold increase in frequency of cysteine (C) in eukaryotes and viruses compared with prokaryotes which is consistent with the literature[25]. The differences in amino acid profiles across proteomes were observed when we analysed proteomes without codon standardisation (Supplementary file 1; F_3,111720_ = 419.57, p < 0.001) as well as for proteomes standardised using amino acid molecular weight, which is known to correlate negatively with amino acid frequency[24] (Supplementary file 1; Domain*Amino acid: F_3,111720_ = 452.12, p < 0.001). These results show that amino acid profiles across proteomes are not conserved in species from distant domains of life.

**Figure 1.**
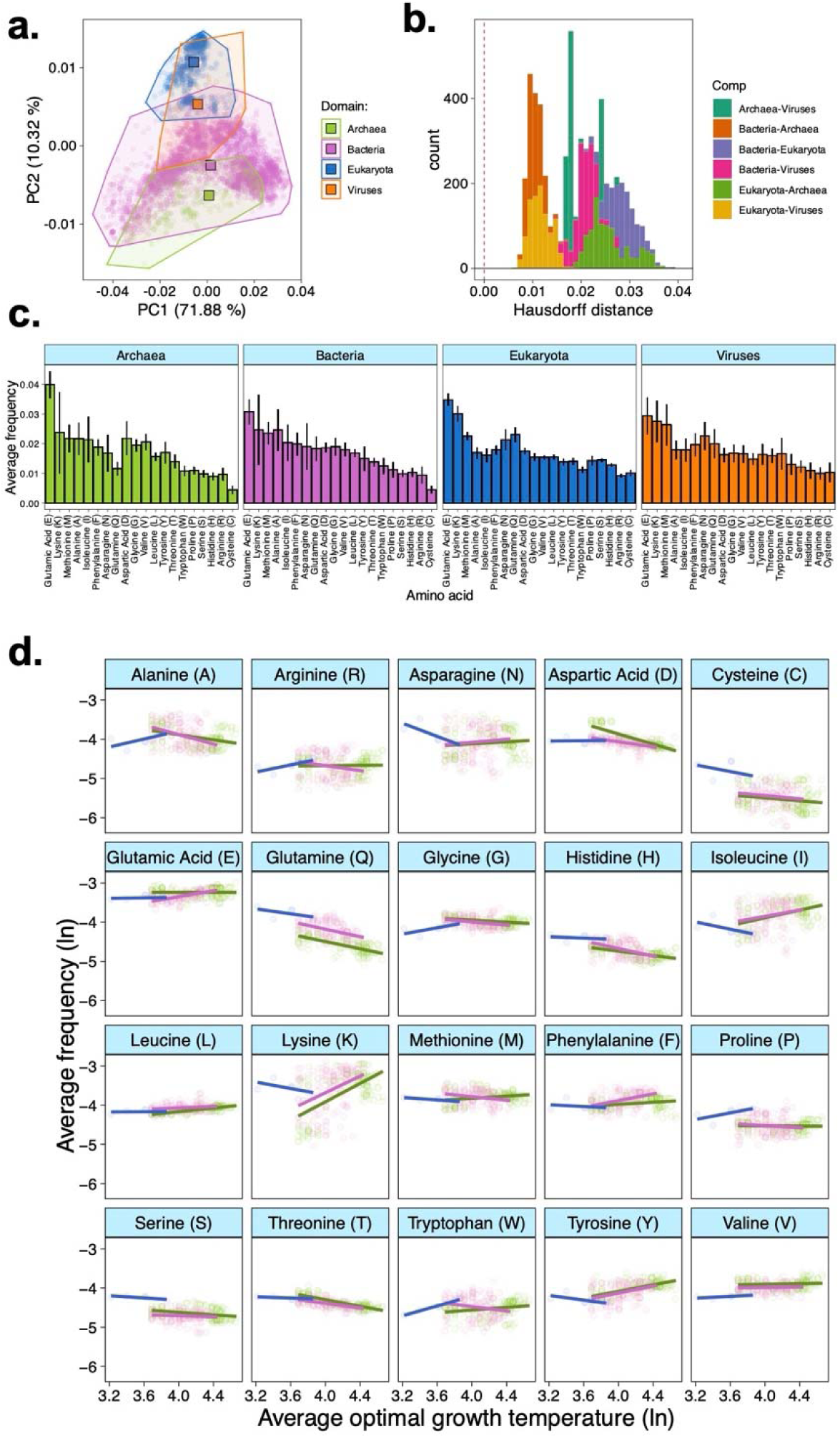
Amino acid frequency across proteomes and the effects of growth temperature. (A) Principal component analysis (PCA) reveals that amino acid usage across genomes in the four domains of life differ. Dots represent the centroids for each of the clusters. (B) Sampling distribution of Hausdorff distances comparing the PCA clusters. Values for the Hausdorff distances equating zero represent no differences between two sets of points. (C) Average amino acid frequencies in proteomes across four domains of life. (D) The relationship between amino acid frequency and optimal growth temperature reveals that environment only has a minor effect on amino acid profiles. In panel (D), axes were ln-transformed.

### Environment weakly explains variation in proteomes

Past studies hypothesised that variation in amino acid profiles were driven by environmental conditions such as optimum growth temperature[5, 14, 15, 26]. Thermophilic proteomes were shown to be under stringent evolutionary constraints[27] which affects amino acid profiles relative to mesophilic proteomes[4, 26] and can lead to highly contrasting amino acid usage profiles[4]. However, these studies were taxonomically biased because they lacked representation of large sets of mesophilic archaea[4, 14, 26]. We incorporated to our proteome dataset information on optimal growth temperature (in °C) for 296 species of Eukaryotes (*n* = 5), Bacteria (*n* = 149) and Achaea (*n* =141) obtained from the ThermoBase database[28] and the supplementary data in[24] (Extended Data 2). We firstly measured how much additional variance was explained when temperature was added as covariate in the model of amino acid frequencies for the 296 species for which growth temperature was available. Temperature had a statistically significant but small contribution to explaining the variance in our model (Likelihood ratio:*χ*^2^-value: 1113.2, p < 0.001; *R^2^* with vs without growth temperature: 0.831 vs 0.797), suggesting that the effects of growth temperature on amino acid profiles were small. This does not contradict previous comparisons of proteomes using pairwise or sophisticated data transformations[4, 14, 26] but it shows that the magnitude of the influence of the environment on the proteome is minor.

Next, we investigated the strength of the relationship between amino acids and growth temperature to disentangle which amino acids, if any, were linked to increasing growth temperatures. For instance, cysteine has been considered an environment-sensing amino acid and is also thermolabile[29]. Our data showed that amino acid frequencies differed with increasing growth temperature among domains of life (Growth Temperature*Domain*Amino acid: F_38,5800_ = 3.005, p < 0.001). This was driven primarily by the an increase in frequency of Alanine (A) and Arginine (R) alongside a strong decrease in frequency of Asparagine (N) and Lysine (K) with increasing growth temperature in eukaryotes but not in archaea or bacteria, a decrease in frequency of Aspartic acid (D) with increasing growth temperature in archaea but not in bacteria and eukaryotes, and an increase in the relative frequency of Glutamic acid (E) and Phenylalanine (F) with increasing growth temperature in bacteria but not archaea or eukaryotes (Fig 1b). There were also statistically significant main effects (Amino acid: F_19,5800_ = 5.526, p < 0.001) and two-way interactions (Growth temperature*Amino acid: F_19,5800_ = 2.384, p < 0.001; Domain*Amino acid: F_38,5800_ = 3.280, p < 0.001) on amino acid frequency. These results confirmed a previous report[26] that increasing growth temperature leads to an overall decrease in frequency of the thermolabile amino acids such as cysteine (C) and glutamine (Q)[5, 30, 31], a pattern which we observed in our data in eukaryotes, archaea and bacteria. Cysteine has also been considered an anomalous amino acid due to lower frequencies than expected by cost models across proteomes[14, 24, 30] although these studies did not directly control for growth temperature which could explain the relatively lower cysteine frequency than expected. Nevertheless, our results show that despite its low frequency, thermolabile amino acids in the proteome negatively correlate with higher optimal growth temperatures. More broadly, our results show that environmental effects in proteomes are minor.

### Few amino acids predominantly appear in first and last ranks

We then ranked amino acids from most to least frequently used in proteomes to investigate their usage frequencies by rank. Our rationale was that amino acid usage by rank could reflect the costs of amino acid usage assuming cost-minimization[4, 32]. Our data shows that the proportion of amino acid by rank were similar across domains of life (Domain*Rank*Amino acid: F_57,771_ = 1.097, p = 0.294), supporting the assumption that amino acid usage by rank is shaped by universal cost-minimization constrains across domains of life. Our data also showed that amino acids were not used uniformly across ranks (Rank*Amino acid: F_19,888_ = 9.376, p <0.001; Fig 2a). This suggested that some amino acids might be differentially prevalent (or even altogether absent) across ranks, which could highlight patterns of amino acid preferential use or avoidance. To test this, we analysed the diversity of amino acids within each rank to assess how many amino acids (raw counts) and their weighted proportions (Shannon-Wiener diversity index) was present across ranks (see eq. 1 in Methods). Under cost-minimization, only few amino acids are expected to be most or least frequently used, leading to a non-linear relationship between our diversity metrics and rank. Our data showed that amino acid diversity by rank varied linearly and non-linearly with rank across domains in both raw counts (Rank*Domain: F_3,68_ = 3.962, p = 0.011; Rank^2^*Domain: F_3,68_ = 3.408, p = 0.022) and Shannon diversity index (Rank*Domain: F_3,68_ = 3.092, p = 0.032; Rank^2^*Domain: F_3,68_ = 6.438, p <0.001). It also confirmed the strong non-linearity of amino acid usage by rank (Counts: Rank^2^: F_1,68_ = 578.117, p <0.001; Shannon: Rank^2:^ F_1,68_ = 425.21, p <0.001), which was not observed for the linear term (Counts: Rank: F_1,68_ = 0.001, p = 0.967; Shannon: Rank: F_1,68_ = 0.987, p = 0.323). These results corroborate our predictions and highlight a novel edge effect where the diversity of amino acids that appeared in ranks 1-2 and 19-20 was lower than the diversity of amino acids in intermediate ranks, an effect observed across all domains of life (Fig 2b). It is unlikely that the edge effect was a statistical artifact because it was observed when the data was analysed without codon standardization or with standardization by amino acid molecular weight (Supplementary File 1) and for proteomes from increasing growth environment (Supplementary File 2).

**Figure 2.**
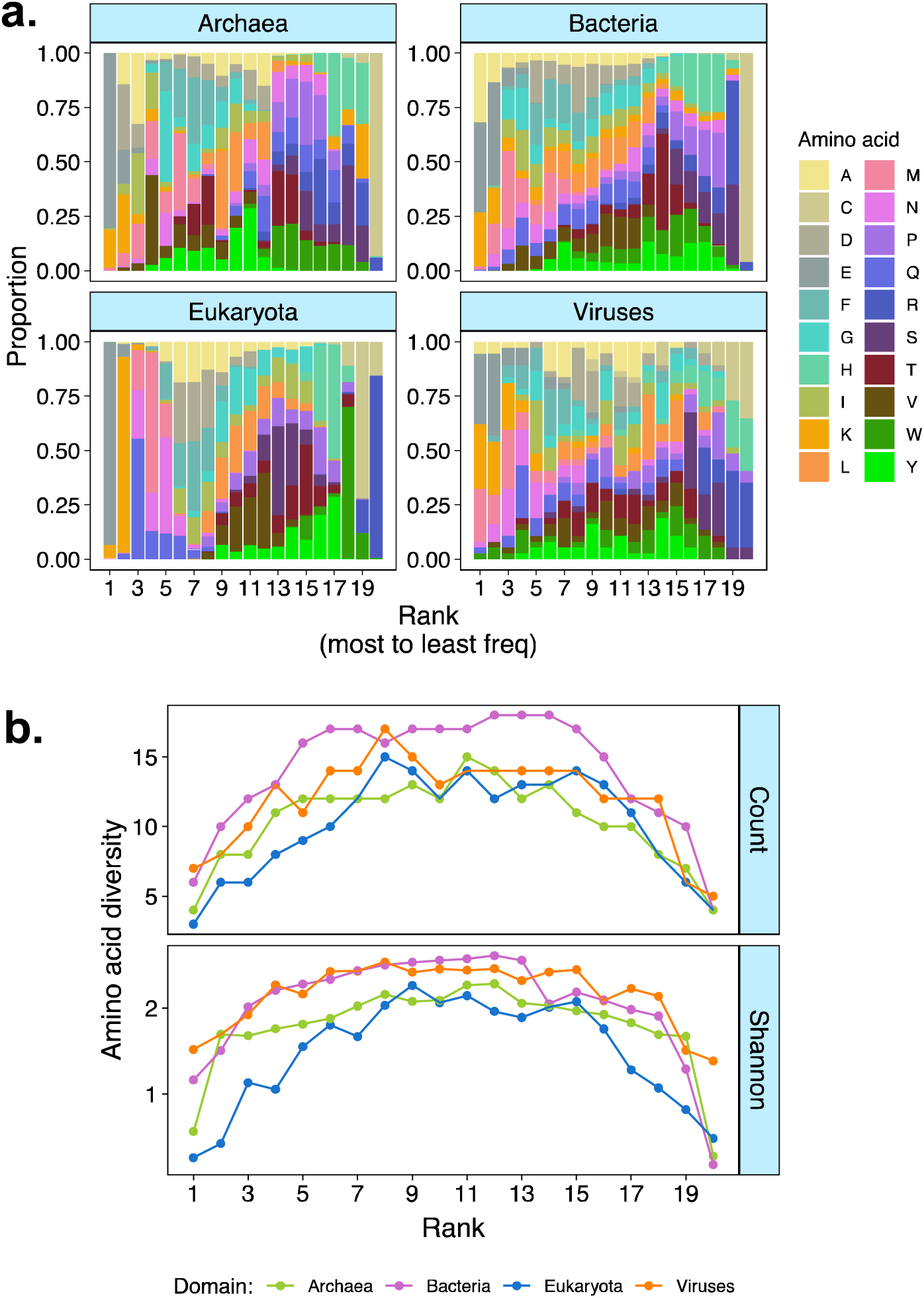
Amino acid diversity decreases in high and low usage ranks. (A) Amino acid proportions by rank across domains of life. (B) Amino acid diversity (Shannon-Wiener index; see Materials and Methods) by rank, showing that diversity decreases at the higher and lower ranks.

### The edge effect is present in the amino acid profiles of secondary structures

The edge effect appears to be ubiquitous and thus, we hypothesised that its cause must also be rooted into fundamental biophysical principles shaping amino acid usage. It is well established that secondary structures such as *α*-helices and *β*-strands are often conserved among protein superfamilies even in distantly related species [33–36]. Moreover, amino acids differ in their propensity to form *α*-helices and *β*-strands[37, 38] which could influence how often they are used in proteomes, depending on their role in secondary structures. Thus, we hypothesised that amino acid frequencies in the proteome reflected their propensity to appear in protein secondary structures, such as *α*-helices and *β*-strands, which could explain the edge effect and why few amino acids appear in most and least frequent ranks in proteomes in all domains of life. To test this, we analysed the amino acid frequency in secondary structures of 40,885 PDB unique entries from 3,512 species across all four domains of life, selected from a subset of structures with low sequence similarity and solved at high-resolution (<3Å) from the PISCES culling database[39, 40]. We first tested whether the average amino acid frequency in the proteome was correlated with the average frequency of the amino acid in *α*-helices and *β*-sheets and found that the average frequency in the proteome and secondary structures were statistically correlated in *α*-helices (Frequency SSE: F_1,72_ = 92.08, p < 0.001) and *β*-sheets (Frequency SSE: F_1,72_ = 8.846, p = 0.003) but differed across domains of life for both secondary structure types (Domain*Frequency SSE*α*: F_3,72_ = 28.279, p <0.001; Domain*Frequency SSE*β*: F_3,72_ = 11.870, p <0.001). This was driven by a weaker positive relationship between amino acid frequencies in the proteome and *α*-helices and a negative relationship between amino acid frequencies in the proteome and *β*-sheets in Viruses compared with other domains of life (Fig 3a).

**Figure 3.**
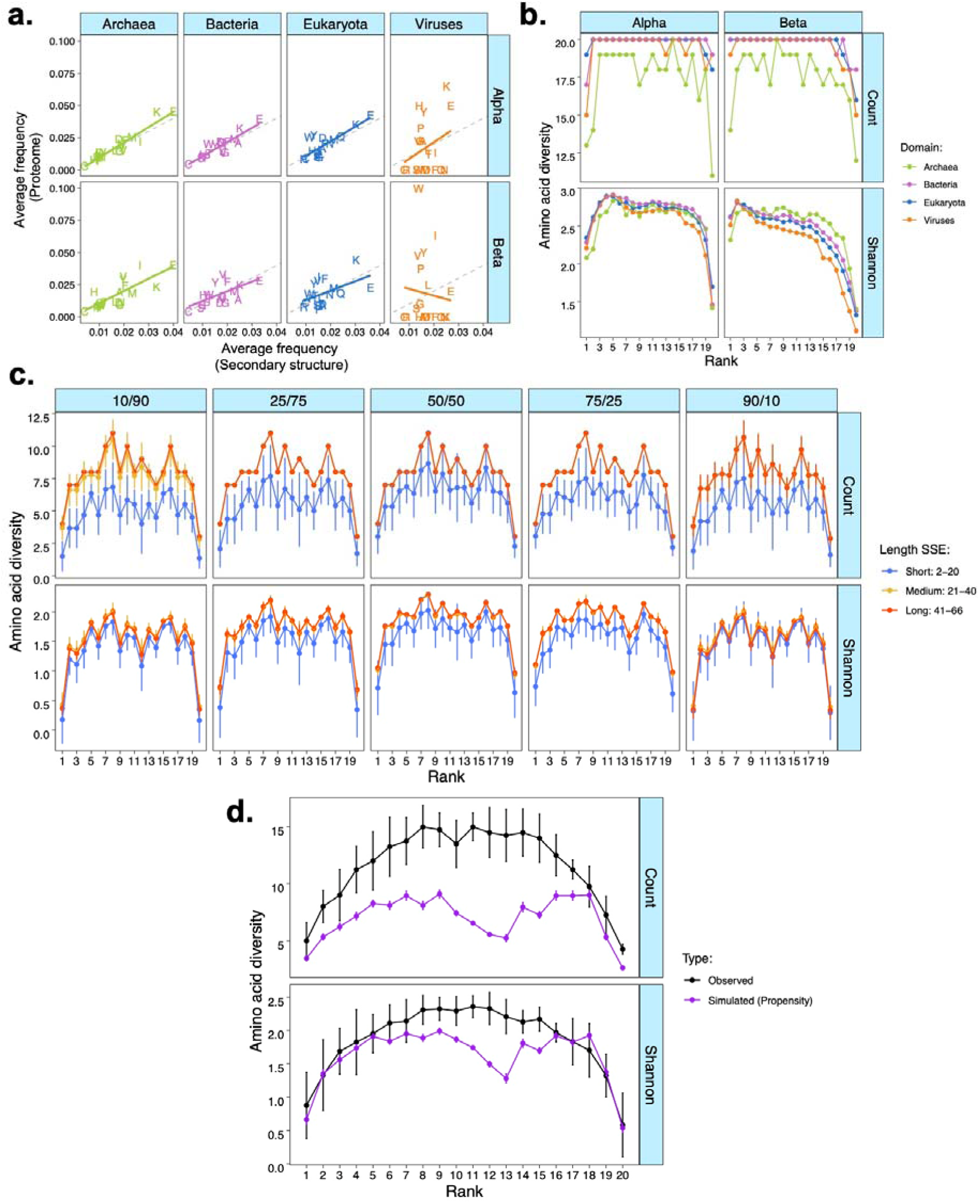
Edge effect is likely driven by conformational properties of amino acids. (A) The relationship between average amino acid frequency in the proteome (y-axis) and on the secondary structures (x-axis). (B) Amino acid diversity by rank within secondary structures. (C) Amino acid diversity by rank of simulated proteins with varying mixtures of *α*-helices/*β*-strands ratio. (D) Comparison between amino acid diversity by rank from the proteome analysis (Observed), simulated data using propensity to form secondary structure from[37] [Simulated (Propensity)].

Next, we hypothesised that the proteome-level edge effect could be an emerging property of the edge effects in secondary structures. We tested this by first ranking amino acids from most to least frequently used in either *α*-helices or *β*-strands for each species and measured the diversity of amino acids in each rank across the domains of life. There was evidence of a non-linear edge effect in both *α*-helices and *β*-strands (Rank^2^: F_1,144_ = 184.833, p <0.001) but this varied depending on secondary structure and domain. For instance, the edge effects on *β*-strands were relatively stronger in higher ranks as opposed to lower ranks, while the edge effects on *α*-helices were more relatively symmetric showing a more characteristic inverted U-shape curve (Rank^2*^SSE: F_1,144_ = 21.719, p <0.001; Fig 3b). The non-linearity pattern of the rank-frequency curve for *β*-strands was less accentuated in viruses which drove a weak but statistically significant interaction between domain and the non-linear effect of rank (Rank^2*^Domain: F_3,144_ = 2.779, p = 0.043; Fig 3b). These results show that the edge effect observed at the proteome-level was also present in the frequency rank of secondary structures, suggesting that the edge effect at the proteome level could be an emerging property of how amino acids form secondary structures.

### The edge effect on amino acid diversity emerges from amino acid conformational properties

Our results for amino acid diversity in secondary structures resembled the distribution of amino acid propensities to form *α*-helices or *β*-strands reported in the classical work by Chou and Fasman (1974)[37], where distributions of amino acid propensities for *α*-helices were relatively symmetric while the propensity for *β*-strands were rightly skewed (Supplementary File 3). This led us to hypothesise that amino acid secondary structure propensities could be the biophysical constrain that gives rise to the edge effect at the proteome level, because amino acids could be differentially selected and used based on their secondary structure propensities. To test whether secondary structure propensity could alone replicate our findings, we simulated 54,400 sequencies of *α*-helices and *β*-strands of varying lengths (from 6 to 66 in increments of 8 residues) where amino acids composition of these simulated secondary structures were selected with probability based only on their propensity to form *α*-helices and *β*-strands as in[37]. This gave us a pool of *α*-helices and *β*-strands with varying amino acid profiles which were representative of their secondary structure propensities. From this pool of simulated secondary structures, we randomly sampled *α*-helices and *β*-strands to assemble 153,450 virtual proteins containing a mixture of these secondary structures. We simulated virtual proteins that were small (2-20 secondary structures), medium (21-40 secondary structures) or large (41-66 secondary structures), each of these with a mixture of secondary structures ranging from proteins that were primarily made of *α*-helices (90%-10%), balanced (50%-50%) or *β*-strands (90%-10%). As expected, the mixture of secondary structures (F_4,8970_ = 178.066, p <0.001) and length (F_2,8970_ = 162.58, p <0.001) of the simulated proteins influenced amino acid rank diversity. However, there was strong evidence that the edge effect could be rescued (Rank^2^: F_1,8970_= 258.30, p <0.001; Fig 3c) independently of length and mixture (Length* Rank^2^: F_2,8970_ = 0.069, p = 0.932; Mixture* Rank^2^: F_4,8970_ = 0.790, p = 0.531; Fig 3c). The edge effect disappeared in simulations where amino acids were drawn with equal probabilities (Supplementary File 4 and Supplementary File 5), supporting that the edge effect is an emerging property of amino acid-specific secondary structure propensities.

We then tested how the simulations compared to our observed data in relation to the edge effect. To do this, we compared the rank-frequency curve from the proteomes in our data base with the curve obtained from the simulations using propensity to form secondary structures from literature[37]. We recapitulated the same edge effect observed in our proteome dataset with our simulation parameterised solely with amino acid secondary structure propensity as in [37]. Both amino acid diversity (Rank^2^*Data type (simulated vs observed): F_1,36_ = 0.172, p = 0.680) and amino acid count showed evidence of edge effects comparable to our observed data (Rank^2^*Data type (simulated vs observed): F_1,54_ = 0.454, p = 0.504; Fig 3d). Simulations had lower raw amino acid counts per rank (F_1,36_ = 28.963, p <0.001) although not lower diversity (F_1,36_ = 3.135, p = 0.085; Fig 3d). These results confirm that amino acid secondary structure propensities could underpin the edge effect on amino acid rank diversity.

### Amino acid usage rank is independent of their evolutionary origin

The evolutionary origin of amino acids into the proteome could influence the frequency in which they are used and therefore, contribute to the edge effect. Two consensus sequencies of amino acid evolutionary chronology exists[41] and we tested whether the average rank of amino acids based on their usage matched their average rank based on their evolutionary chronology. There was no evidence that average amino acid rank from frequency correlated with average rank from evolutionary chronology across domains of life for neither (Raw: F_3, 72_ = 0.405, p = 0.749; Filtered: F_3, 72_ = 0.390, p = 0.759; Table 1). These results show that amino acid rank usage is determined by functional constrains on their use above evolutionary chronology.

**Table 1.** Relationship between average rank from amino acid usage and average rank from evolutionary chronology as in[41]. ‘Raw’: unfiltered average chronology ranks from the 40 criteria used in Trifonov (2000)[41]. ‘Filtered’: filtered average chronology ranks, accounting for correlation between the 40 selection criteria as in Trifonov (2000)[41]. Slopes ± 95% Confidence intervals) reported. P-values estimated from F-statistics. (attached)

## Conclusion

We showed that a few amino acids are most and least frequently used in proteomes across domains of life and that this effect was driven by amino acid secondary structure propensities. This has implications to our understanding of the dynamics underpinning protein structure evolution and to our estimates of amino acid substitution rate in matrices used for amino acid sequence alignments to probe deep evolutionary history[42]. This is because the edge effect, which was pervasive across domains of life, indicates that few amino acids might be consistently more, or less, frequent in the proteome which may influence the substitution rates estimated in these matrices. We also showed that, despite the exceptions related to the edge effect, the overall amino acid usage profile of the proteomes varied among domains contradicting recent hypotheses[15, 16]. Our results also showed that environmental conditions as measured by optimal growth temperature had a significant but minor effect on the amino acid frequency, including of thermolabile amino acids. Previous studies predicted a larger effect of optimal growth temperature on proteomes[4]. This discrepancy is likely because our dataset has sampled a higher proportion of mesophilic species, providing a better resolution for the effects of optimal growth temperature on amino acid frequency across proteomes on a continuous scale. Collectively, our findings give new insights into the relationship between structure and function of amino acid and proteins, highlighting a novel universal constrain shaping amino acid usage across proteomes. Proteomes contain critical knowledge about genome and protein evolution[4, 5] and can inform new ways to probe deep evolutionary histories between proteins with potential applications to protein engineering strategies in the era of synthetic biology[43, 44].

## Materials and Methods

### Amino acid profiles and growth temperature

All analyses and simulations were conducted in R 4.3.2[45]. NCBI data was accessed and retrieved on or before January 2^nd^ 2024. Translated coding sequences were downloaded from FTP servers and fasta files were processed in the statistical software R version 4.3.2[45] to estimate amino acid profiles. The list of specie and their NCBI accession number is given in Extended Data 1. Only reference sequence (RefSeq) genomes were considered to ensure maximum coverage and annotation accuracy. For consistency of analysis across domains, we included proteomes from hosts without intracellular organelles. Because of the positive correlation between the number of redundant codons and the frequency of amino acids, we standardised our amino acid profiles, dividing amino acid frequency by the number of redundant codons. To confirm that our findings, particularly related to the edge effect, were not due to the standardization, we also analysed our amino acid profile data using no-standardization (Supplementary File 1) or standardization by amino acid molecular weight (Supplementary File 2). After standardization by codon, there was no statistical relationship between the natural log-frequency and natural log-ATP costs controlled by codon redundancies based on the amino acid costs reported in[46] (F_1,72_ = 1.349, p = 0.249, Supplementary File 5). Average optimum growth temperature was retrieved from the ThermoBase database[28] and consolidates with supplementary data for eukaryotes from the supplementary material in[24]. ThermoBase was accessed on March 1^st^ 2024. The final dataset of 296 species of which 142 Archaea and 149 Bacteria and 5 Eukaryotes (Extended Data 2) were used. We validated our calculations of amino acid frequencies using an independent dataset (i.e. RACCOON dataset[47]) which contained 26 species that were also present in our dataset. There were no statistically significant differences between the estimates of amino acid frequency between our dataset and the RACCOON dataset for the 26 species (*t*-value = 1.295, df = 519, p = 0.196) and the edge effect found here was also observed in the RACCOON dataset (Supplementary File 5). For our analysis of amino acid frequency in secondary structures, we downloaded the PDB structural information using the ‘bio3d’ package[48] of 40,885 PDB unique entries from 3,512 species across all four domains of life, selected from a subset of the PISCES PDB culling database[39, 40]; PDB entry IDs are given in the Extended Data 3.

### Statistical analysis

Plots were made using the ‘ggplot2’ package[49] and mixed linear regressions were fitted using the ‘lme4’ and ‘lmerTest’ packages[50, 51]. To test if amino acid profiles differed across domains of life, we adopted two approaches. First, we ran a principal component analysis (PCA) using the ‘prcomp’ inbuilt R function which revealed the amino acid usage profile of each domain of life. To ascertain whether these profiles were different, we compared the clusters using a bootstrapping approach with 1000 replicates and calculating the Hausdorff distance between bootstrapping samples. Hausdorff distances applied to biological data is explained in details here[52] but briefly, Hausdorff distance is a distance metric used to compare two sets. When the distance value is zero, the two sets are identical. For inference purposes, we assumed that, since all pairwise comparisons resulted in Hausdorff distance greater than zero, the amino acid profiles shown in the PCA analysis were different. We also fitted a linear mixed model using amino acid frequency as dependent variable, amino acid type and domain of life (and their interaction) as fixed effects, and species as random effects to account for multiple measurements of amino acids for each species. These two approaches revealed similar patterns. A similar mixed model was fitted to analyse the effects of growth temperature but incorporating average optimal growth temperature and the three-way interactions with amino acid type and domain of life. In all models, amino acid frequency was log-transformed to improve fit of the data. Likelihood ratio test to evaluate the increase in explanatory power by adding growth temperature was done by fitting two models, with and without growth temperature, and comparing using the ‘anova’ function of the ‘stats’ package[53], as well as the ‘r-squaredGLMM’ function of the ‘MuMIn’ package[54] to obtain conditional R^2^ for our mixed models. Proportion of curve classes within each domain of life was done using the ‘chisq.test’ function of the ‘stats’ package. We fitted a binomial generalized linear model with *quasi* errors using the ‘glm’ function to analyse the proportion of amino acid per rank as response variable and the three-way interactions between rank, amino acid, and domain of life as independent variables. For analysis of the relationship between amino acid average frequency in the proteome and on secondary structures, we used a general linear model with average frequency in the proteome as response variable and the interaction between average amino acid frequencies in the secondary structures and domain as independent variables. We estimated amino acid diversity per rank as the count of different amino acids per rank as well as the Shannon-Wiener diversity index per rank using the ‘tabula’ package[55] which estimates the index using the following equation:

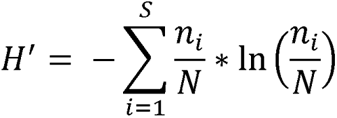

Where *S* represents the total number of amino acids *i* by rank and *ln* is the natural logarithm.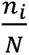 represents the proportion of each amino acid *i* by rank[55] (see also[56]). Because we were interested in edge effects, which are non-linear (i.e. inverted U-shape), we fitted general linear models using either diversity metric as dependent variable, the non-linear effects of rank and its interaction with domain for observed data or the non-linear effects of rank and their interaction with sequence length, mixture of secondary structures, and their three-way interactions for our simulations. To compare whether the edge effect was rescued in our simulations, we fitted a general linear model with either diversity metric as dependent variable, the non-linear effects of rank and its interaction with data type (i.e. simulation from propensity (“Propensity”, simulation using PDB data (“Data”) and observed proteome-level data; Fig 3d). We also regressed average rank estimated by two methods of evolutionary origins of amino acids in the genetic code from [41] with the weighted average rank of each amino acid from our proteome analysis.

## Declarations

### Ethics approval and consent to participate

Not applicable

### Consent for publication

Not applicable

### Availability of data and materials

All data are provided as supplementary material.

### Competing interests

The authors declare that they have no competing interests

### Funding

JM is supported by the Biotechnology and Biological Sciences Research Council (BBSRC: BB/V015249/1)

### Authors’ contributions

JM and ZP conceptualised the study. JM and ZP downloaded the data. JM analysed the data and created the figures. JM and ZP wrote and edited the manuscript.

## Acknowledgements

We thank Prof Thomas Richards for advice on the early stages of the analysis.

## Authors’ information

JM is an early career researcher and senior lecturer at the Institute of Mathematics at the University of Aberdeen (UK). He is a biologist who works at the interface of ecology, computational biology, and mathematics. He is also an honorary professorship at the Federal University of Paraná (Brazil), where he graduated with a BSc in Biological Sciences in 2013. He graduated in 2016 with a DPhil in Zoology by the University of Oxford. ZP is an early career researcher at the transition to her first postdoctoral role at Lund University (Sweden). She graduated in 2022 from Linköping University (Sweden), before entering maternity leave for over a year.

## Supplementary files

**Extended Data 1.** Amino acid profiles for 5,590 species.

**Extended Data 2.** Growth temperature for 296 species of Bacteria and Archaea for which amino acid profiles were estimated.

**Extended Data 3.** PDB identification numbers for the structures analysed in this study.

**Supplementary File 1.** (a) Amino acid profiles differ between domains of life as shown by principal component analysis and the Hausdorff distance in non-standardised and molecular weight-standardised data. (b) Edge effect present in amino acid profiles for non-standardised and standardised by molecular weight models.

**Supplementary File 2.** Proportions of amino acid by rank for proteomes of species with optimal growth temperature from 0 to 70oC and from 70°C to 110°C (extremophiles).

**Supplementary File 3.** Density plot of the distributions of amino acid propensities for *α*-helices (symmetric) and *β*-strands (right-skewed).

**Supplementary File 4.** The edge effect disappears in simulations where amino acids were sampled with equal (0.05) probabilities, confirming that differences in secondary structure propensities drive the edge effect.

**Supplementary File 5.** (a) Differences in estimated amino acid frequency between our data and the RACCOON dataset. (b-c) The edge effect described here was also present in the RACCOON dataset. (d) The relationship between amino acid frequency and ATP/time costs from[46].

## Notes

### Competing Interest Statement

The authors have declared no competing interest.

