## Supplementary material for "Differential amino acid usage leads to ubiquitous edge effect in proteomes across domains of life that can be explained by amino acid secondary structure propensities": Fig S1

### Slide 1
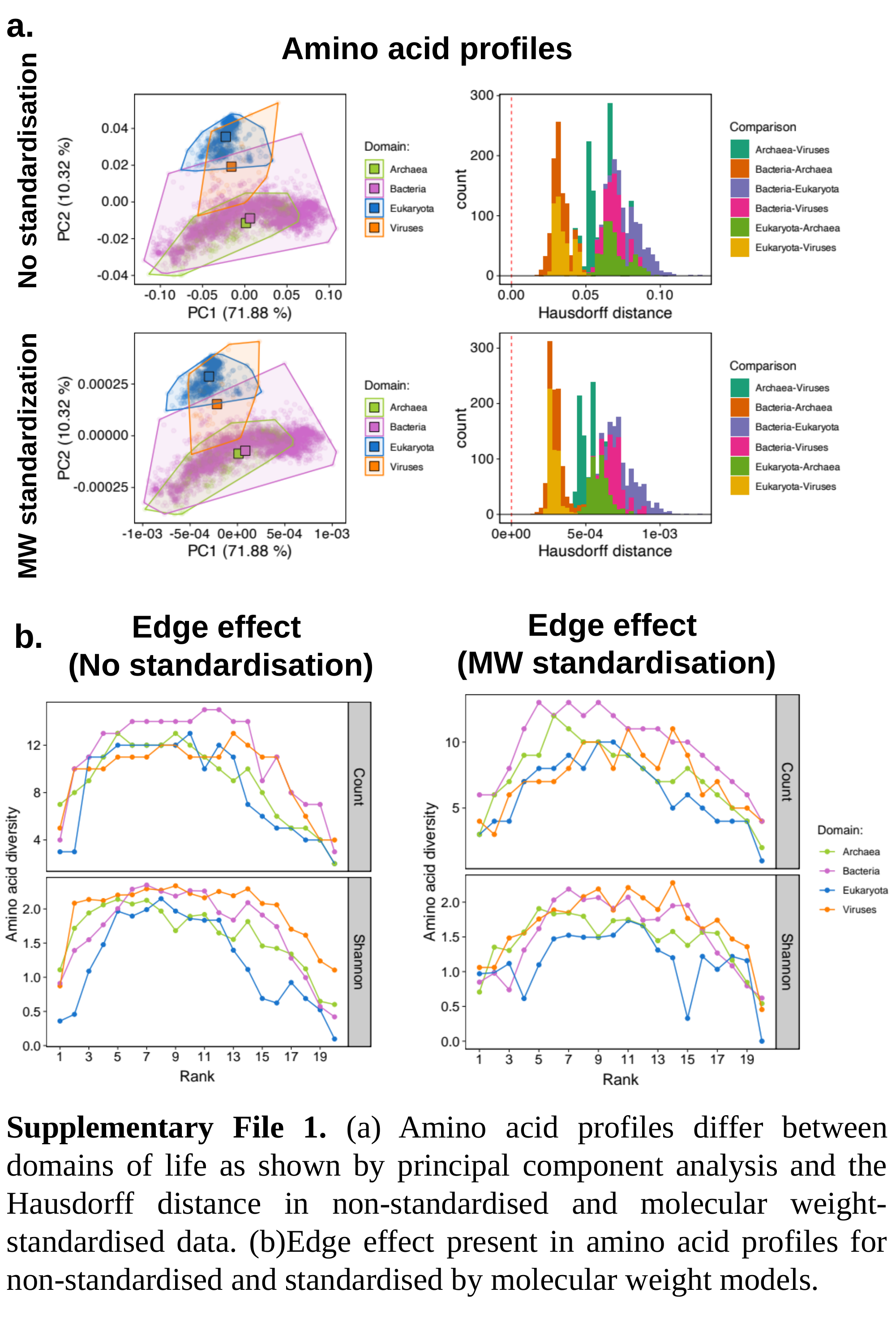

a.
Amino acid profiles
 MW standardization No standardisation
Edge effect
(MW standardisation)
Edge effect
(No standardisation)
b.
Supplementary File 1. (a) Amino acid profiles differ between domains of life as shown by principal component analysis and the Hausdorff distance in non-standardised and molecular weight-standardised data. (b)Edge effect present in amino acid profiles for non-standardised and standardised by molecular weight models.
