## Supplementary material for "Differential amino acid usage leads to ubiquitous edge effect in proteomes across domains of life that can be explained by amino acid secondary structure propensities": Fig S2-S4

### Slide 1
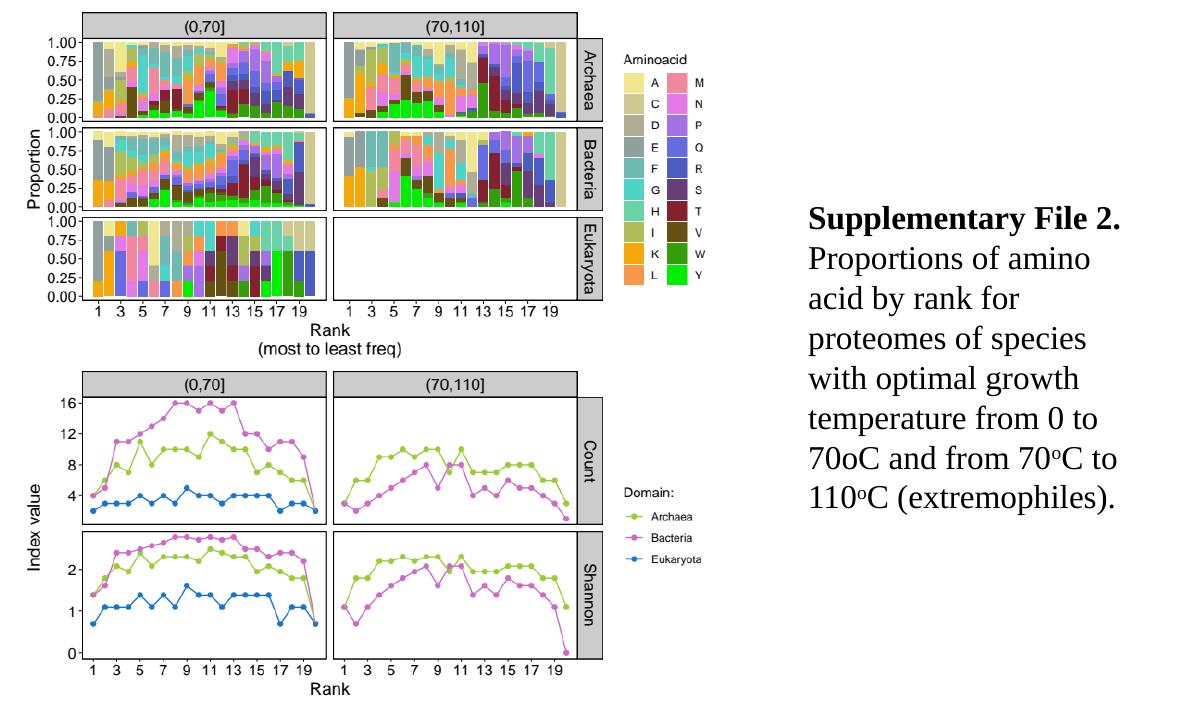

Supplementary File 2. Proportions of amino acid by rank for proteomes of species with optimal growth temperature from 0 to 70oC and from 70^o^C to 110^o^C (extremophiles).

### Slide 2
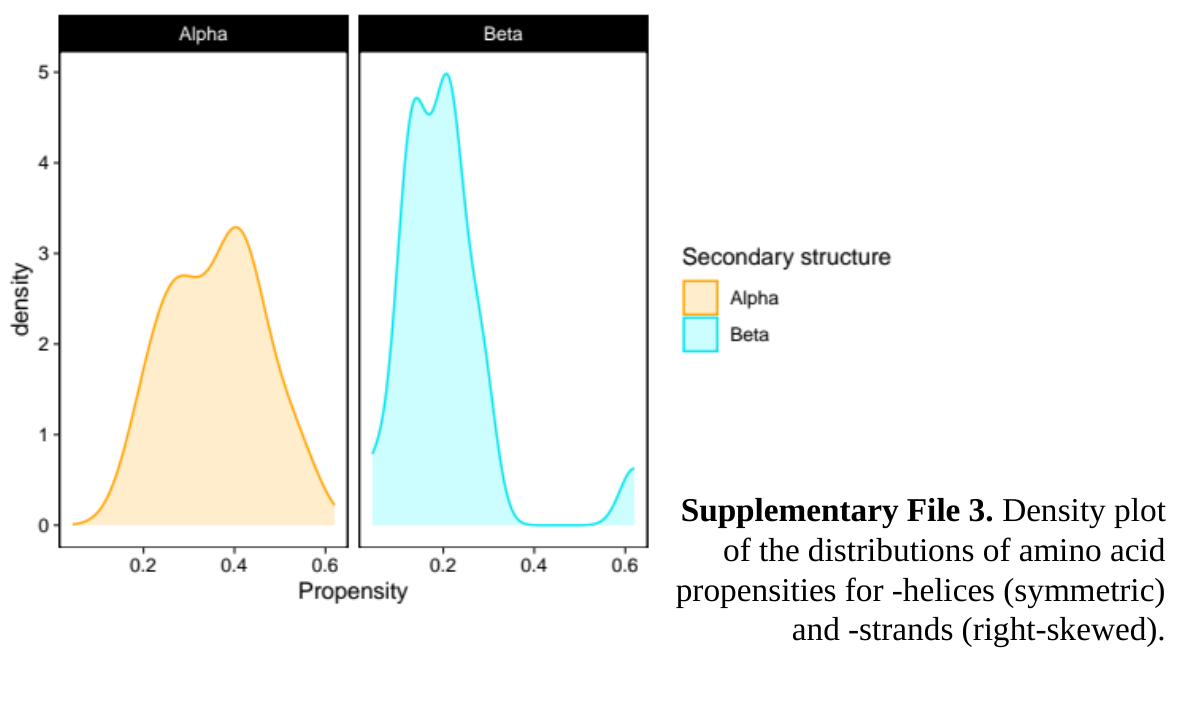

### Slide 3
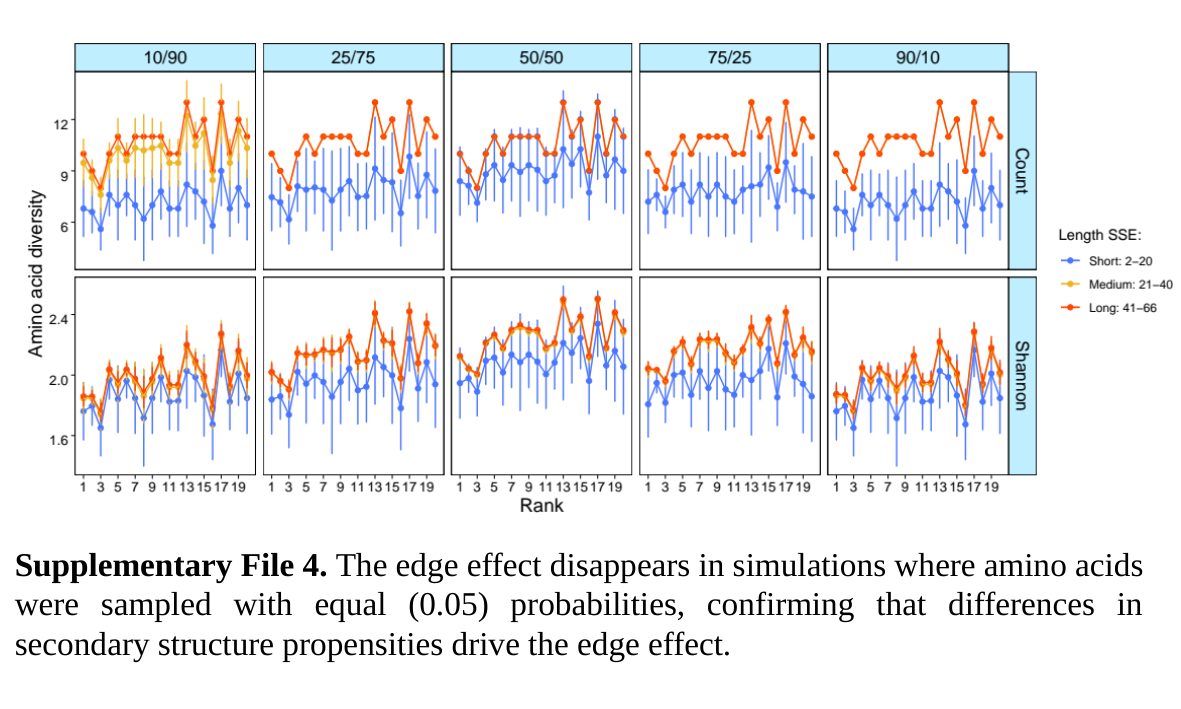

Supplementary File 4. The edge effect disappears in simulations where amino acids were sampled with equal (0.05) probabilities, confirming that differences in secondary structure propensities drive the edge effect.
