## Supplementary material for "Differential amino acid usage leads to ubiquitous edge effect in proteomes across domains of life that can be explained by amino acid secondary structure propensities": Fig S5

### Slide 1
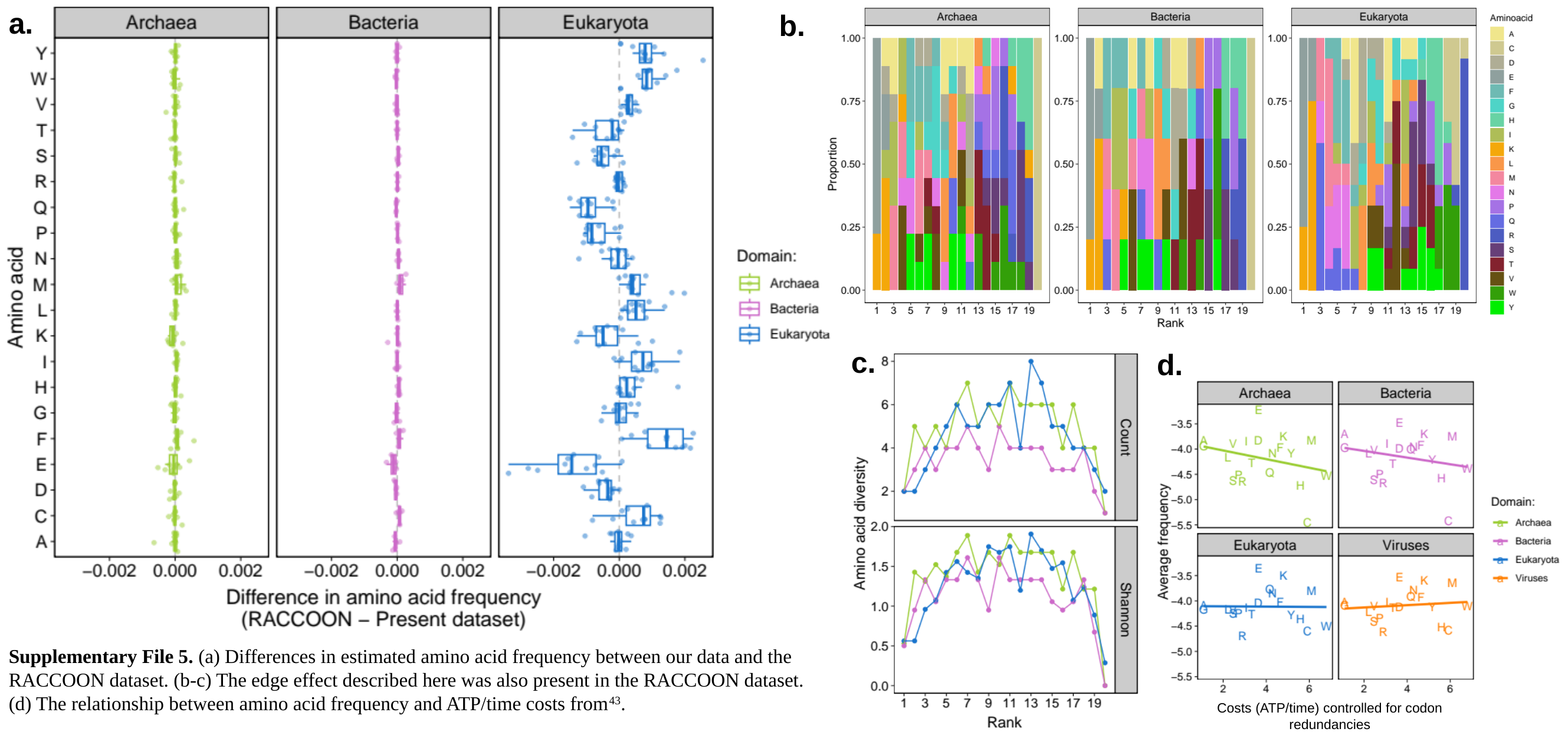

a.
b.
c.
d.
Supplementary File 5. (a) Differences in estimated amino acid frequency between our data and the RACCOON dataset. (b-c) The edge effect described here was also present in the RACCOON dataset. (d) The relationship between amino acid frequency and ATP/time costs from^43^.
Costs (ATP/time) controlled for codon redundancies
